# A Counterfactual Framework for Directional Cell–Cell Interaction Analysis in Spatial Transcriptomics

**DOI:** 10.64898/2026.04.05.716510

**Authors:** Humaira Anzum, Veena Kochat, Suresh Satpati, Md Ishtyaq Mahmud, Jagan Mohan Reddy Dwarampudi, Pooja Shukla, Milind Javle, Lawrence Kwong, Kunal Rai, Tania Banerjee

## Abstract

Understanding how neighboring cells influence cellular states is central to spatial transcriptomics, yet most existing methods rely on correlation or predefined ligand–receptor (LR) pairs and do not explicitly test directionality. We introduce a counterfactual, intervention-based framework for inferring directional cell–cell influence that is LR-agnostic and tests sender specificity.

A neighborhood-conditioned graph model predicts receiver cell state from local spatial context. Directional influence is quantified by counterfactually replacing neighbors of a candidate sender type and measuring the resulting displacement in predicted receiver state. We define a Counterfactual Directionality Score (CDS) that quantifies directional influence, and compute pair-level CDS by aggregating across receiver cells and test cores for each ordered sender–receiver pair.

Applied to Xenium cholangiocarcinoma tissue microarrays (38 cores), the framework identified reproducible, asymmetric interactions between tumor, immune, and stromal compartments, most prominently Tumor-EMT → Macrophage (CDS = 0.0828) and Fibroblast → Macrophage (CDS = 0.0582). Effects exceeded label-permutation and spatial-shuffle null models (*p <* 0.001, FDR-controlled) and remained stable under core-level bootstrap resampling. Inferred directional strengths correlated strongly with matched LR scores (*r* = 0.758, *p* = 0.0027), supporting biological concordance.

These results demonstrate counterfactual testing as a statistically rigorous and scalable approach for directional cell–cell communication analysis in spatial transcriptomics.

## 1 Introduction

Spatial transcriptomics measures gene expression while preserving tissue architecture, enabling analysis of cell–cell interactions in situ [4]. In oncology and immunology, understanding how specific cell types influence neighboring cells is central to tumor progression, immune modulation, and therapeutic response [11,7]. However, inferring *directional* cell–cell influence from single-snapshot spatial data remains challenging [10].

Existing computational approaches to cell–cell communication (CCC) are largely correlational. Ligand–receptor (LR)-based methods, including CellPhoneDB [5], CellChat [6], and NicheNet [1], test curated interaction pairs between annotated populations. While interpretable, they depend on prior knowledge and do not explicitly test directionality [8]. Spatial toolkits and transport-based models (e.g., Squidpy [9], Giotto [3], SpaOTsc [3], COMMOT [2]) incorporate spatial structure, and predictive frameworks such as GNNs and transformer-based models (e.g., GITIII [12]) capture complex dependencies. Nevertheless, influence is inferred through predictive attribution rather than explicit intervention.

We instead adopt a counterfactual formulation of directional influence. A neighborhood-conditioned graph model predicts receiver cell state from local spatial context, after which controlled perturbations of sender composition quantify the resulting displacement in predicted receiver state. This enables directional inference through interventional testing rather than association.

### Contributions

(1) We introduce a counterfactual framework for directional cell–cell influence in spatial transcriptomics without predefined LR priors. (2) We define a Counterfactual Directionality Score (CDS) with complementary sender type-swap and within-type shuffle operators to disentangle cell-type identity from intra-type heterogeneity. (3) We establish a rigorous evaluation protocol using matched permutation null models with FDR correction and strict core-level data separation. (4) We enhance interpretability through functional program projection and ligand–receptor concordance analysis.

## 2 Proposed Methodology

### 2.1 Overview

We introduce a counterfactual framework to quantify directional cell–cell communication in spatial transcriptomics. Cell state vectors are derived by subtracting cell-type means, then a neural network (the *Neighbor Influence Model*) is trained to predict a receiver cell’s state from its spatially weighted neighbors. We measured the directional influence between cell types by perturbing sender composition within neighborhoods and quantifying changes in predicted receiver state across independent tissue microarray (TMA) cores.

### 2.2 Cell State and Spatial Context Modeling

Let *x*_*i*_ ∈ ℝ^*G*^ (*G* = 480) denote the expression vector of cell *i*. To isolate functional state from cell-type identity, we compute a residualized state *s*_*i*_ = *x*_*i*_ − *µ*_*t*(*i*)_, where *t*(*i*) is the cell type and *µ*_*t*_ its mean expression. Residual values are clipped to [−5, 5] and projected to 50 principal components, yielding a denoised state representation *z*_*i*_ ∈ ℝ^50^.

Each TMA core is treated as an independent biological unit. For every cell *i*, the *k* = 20 nearest spatial neighbors are identified using Euclidean distance in tissue coordinates. Neighbor states are aggregated via distance-weighted softmax pooling:

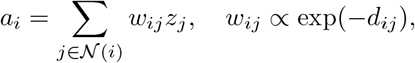

producing a neighborhood context vector *a*_*i*_ ∈ ℝ^50^.

We train a *Neighbor Influence Model* to predict receiver state *z*_*i*_ from neighborhood context *a*_*i*_ conditioned on receiver cell type *t*(*i*). The model comprises an input batch normalization layer followed by residual hidden layers with layer normalization. Receiver cell type is provided as an index tensor (`recv_type_idx`) and mapped to a learnable embedding via an `nn.Embedding` layer. This embedding is linearly projected and passed through a sigmoid to generate a multiplicative gate, which modulates the hidden representation element-wise, enabling type-specific processing of spatial context. The network is trained using Huber loss (*δ* = 1.0) between predicted and observed cell states.

Cores are partitioned into training (25), validation (3), and held-out test (10) sets, with counterfactual analyses performed exclusively on test cores. Optimization uses AdamW with early stopping based on validation loss.

### 2.3 Counterfactual Directionality Score (CDS)

For a sender type *S* and receiver type *R*, we quantify directional influence by comparing model predictions under observed versus counterfactual neighborhoods. For each receiver cell *i* of type *R* in test cores, we define the vector difference:

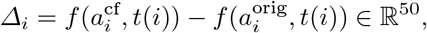

where 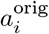 uses the original neighbors and 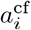 is computed from a counterfactual neighborhood. Two complementary counterfactual operators are applied:

#### Type-swap operator

Neighboring *S* cells are replaced with non-*S* cells drawn from the same core, preserving distance bin structure where possible. This tests whether the presence of sender type *S* causally influences receiver *R*.

#### Within-type shuffle operator

Neighboring *S* cells are replaced with *other* cells of the same type *S* from the same core, preserving sender identity but disrupting specific sender states. This operator tests dependence on sender cell-state heterogeneity.

For each receiver cell *i*, we define the CDS as the L1 displacement

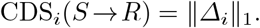

The core-level CDS was calculated by averaging receiver-cell CDS values within each core. Furthermore, the pair-level CDS was computed by averaging core-level CDS values across the 10 held-out test cores.

Signed mean deltas were examined separately to characterize the direction of receiver-state shifts (activation vs. suppression).

### 2.4 Program-Level Projection

To enhance biological interpretability, gene-level changes are projected onto predefined functional programs (e.g., EMT, macrophage activation, T-cell exhaustion). For a given functional program *P*, we compute the program-level delta for cell *i* by first mapping the counterfactual state difference back to gene expression space using the inverse PCA transformation, and then averaging the resulting gene-level deltas across all genes in the program. Program-level CDS are aggregated identically to gene-level CDS. This provides insight into which biological processes are most affected by a given sender–receiver interaction.

### 2.5 Statistical Significance

Statistical significance was assessed using three matched null models (each with *B* = 200 permutations) preserving complementary aspects of biological structure: (i) label permutation within cores (tests dependence on cell-type annotation), (ii) distance-bin neighbor shuffling followed by type-swap (tests spatial specificity), and (iii) sender-agnostic replacement (tests sender identity versus generic occupancy).

Empirical p-values were computed as 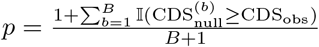, with Benjamini–Hochberg correction applied across all sender–receiver pairs to control the false discovery rate (FDR) at *α* = 0.05.

### 2.6 Cross-Core Uncertainty

Because TMA cores represent independent biological replicates, uncertainty is quantified using block bootstrap at the core level. For *R* = 1000 iterations, test cores are sampled with replacement and the mean CDS recomputed. The bootstrap distribution provides the bootstrap mean and 95% confidence intervals (2.5th and 97.5th percentiles).

### 2.7 Ligand–Receptor Concordance Validation

For orthogonal biological validation, we assess whether genes driving significant CDS values correspond to known ligand–receptor interactions between sender and receiver types. For each significant pair, we identify genes with largest absolute mean delta and query CellChatDB for interactions where the ligand is expressed in the sender type and the receptor in the receiver type. Only pairs with mean expression *>* 0.05 in both cell types are retained. The ligand–receptor score is defined as the geometric mean of sender ligand expression and receiver receptor expression. Concordance was quantified by computing the Spearman rank correlation between pair-level CDS values and corresponding ligand–receptor (LR) scores across all evaluated sender–receiver pairs.

## 3 Experiments

### 3.1 Dataset and Evaluation Protocol

We evaluated the proposed counterfactual framework on Xenium spatial transcriptomics data from human cholangiocarcinoma tissue micro-arrays (TMA).

After doing quality control, we selected 38 independent TMA cores. We then partitioned the cores into 25 training, 3 validation, and 10 held-out test cores. We exclusively computed all reported CDS values, null model analyses, and biological interpretations on the held-out test set. CDS values were first calculated within each core and then aggregated and the uncertainty was estimated using block bootstrap resampling at the core level (B=1000).

### 3.2 Spatial Model Validation

To verify that our Neighbor Influence (NI) model leverages spatial information rather than learning cell-type-specific expression, we performed a controlled validation on held-out test cores comparing three models: (i) true-NI (authentic spatial neighbors); (ii) shuffled-NI (disrupted neighbor-receiver correspondences); and (iii) a receiver-only MLP baseline (predicts cell state solely from cell type).

Figure 1 shows that true-NI consistently achieves lower prediction error than both baselines. In panel A, it can be seen that the mean squared error (MSE) averaged across 10 held-out cores is lowest for true-NI (0.849), compared to MLP (1.032) and shuffled-NI (0.964). The error bars indicate ± 1 SD. Panel B displays per-core paired comparisons. So for nearly every core, true-NI yields reduced error relative to both baselines, with boxplots confirming downward shift in both median and variance. Paired *t*-tests confirm significance (*p <* 0.001), which demonstrates that authentic spatial neighborhoods provide measurable predictive benefit.

**Fig. 1.**
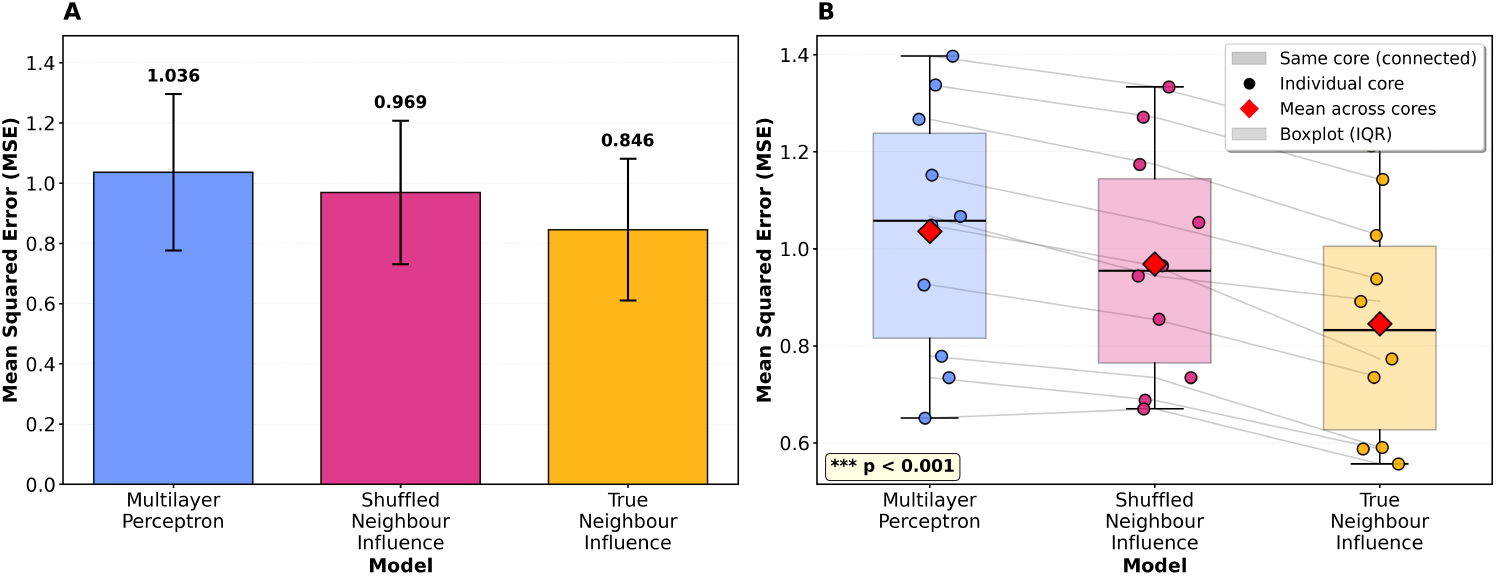
Performance comparison across held-out test cores. (**A**) Cohort-level MSE with *±*1 SD error bars. (**B**) Per-core paired errors with boxplots and mean markers.

**Fig. 2.**
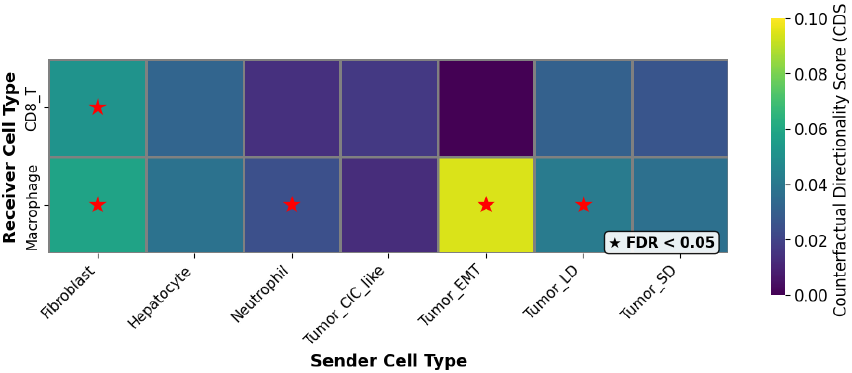
CDS under type-swap analysis. Red stars denote FDR *<* 0.05.

### 3.3 Directional Influence of Sender Cell Types

We evaluated directional influence using a sender type-swap counterfactual operator. For each sender–receiver pair, we replaced neighboring sender cells with non-sender cells of a different type from the same core, preserving all other micro-environmental aspects. The resulting shifts in predicted gene expression were aggregated per core as a CDS. Averaging across the 10 held-out test cores yields the pair-level CDS for each sender–receiver pair. Higher CDS indicates stronger directional influence of the sender on the receiver.

Several interactions exhibited a statistically significant directional influence. For instance, *Fibroblast* → *Macrophage* showed a robust effect (CDS = 0.0582, 95% CI [0.0459, 0.0719], FDR *<* 0.05), while *Tumor-EMT* → *Macrophage* (CDS = 0.0828, 95% CI [0.0811, 0.0839], FDR *<* 0.05) represented the largest directional perturbation detected across all pairs.

### 3.4 Biological Interpretation of Directional Effects

Gene-level counterfactual deltas were projected onto predefined biological programs to assess coherent functional shifts (Figure 3). Tumor-EMT → Macrophage interactions showed significant suppression of Angiogenesis and Macrophage Activation programs, whereas other pairs exhibited weaker or selective effects, indicating program-specific modulation. Sender–receiver pairs showed coherent biological patterns (Figure 4). Tumor-EMT and Tumor-LD reprogrammed macrophage states, while Fibroblast → CD8 T cell interactions suggested stromal suppression of cytotoxic programs. Ligand–receptor concordance analysis using CellChatDB (Figure 4) identified organized interaction structure, with receptors such as *CD44, CCR7*, and *CXCR4* acting as central nodes. This supports interpretation of the directional effects as coordinated signaling modules rather than isolated gene changes (see Section Ligand–Receptor Concordance Validation).

**Fig. 3.**
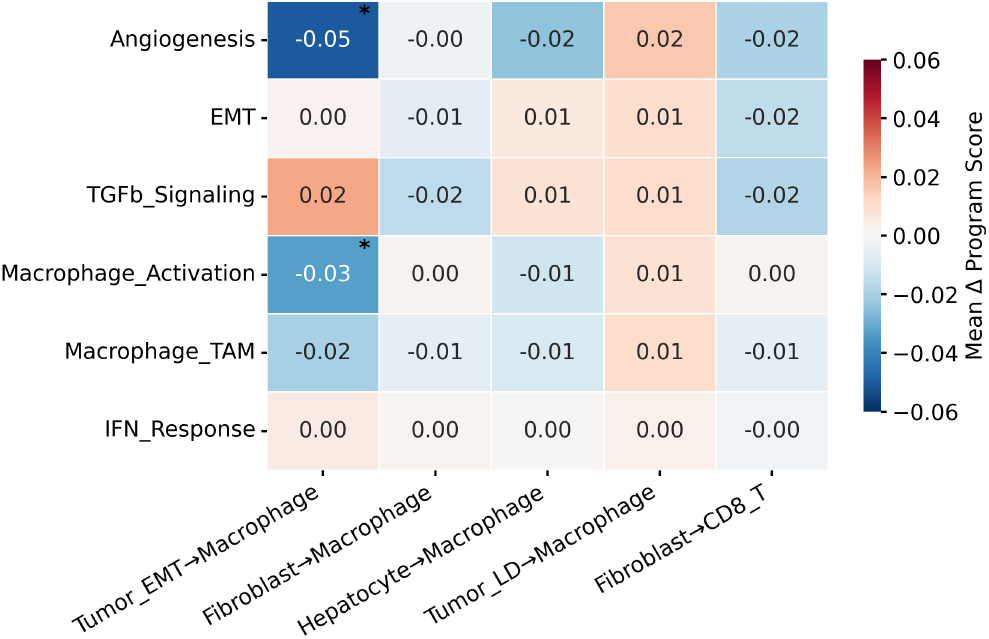
Program-level mean signed counterfactual deltas (asterisk: FDR *<* 0.05).

**Fig. 4.**
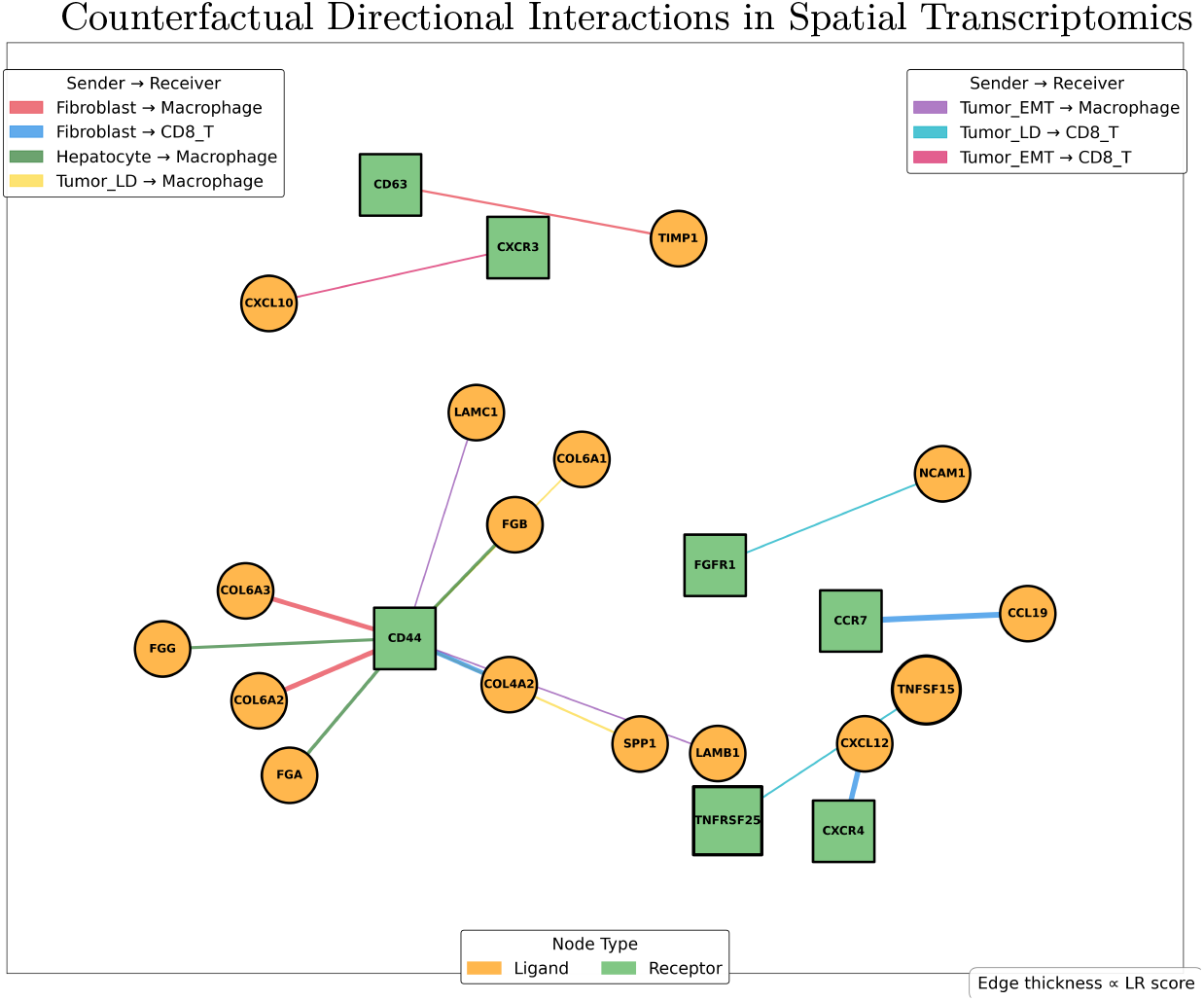
Ligand–receptor interaction network across sender–receiver pairs.

### 3.5 Robustness Analysis

We evaluated whether the observed effects exceed those expected under randomized spatial structure using matched null models (Fig. 5). Under the type-swap operator (Fig. 5A), the Fibroblast → Macrophage CDS (0.0582) lies in the extreme tail of the label-permutation, neighbor-shuffle, and sender-agnostic null distributions (all *p <* 0.001, FDR-significant), demonstrating robust directional influence that is specific to the true spatial and sender-type configuration.

**Fig. 5.**
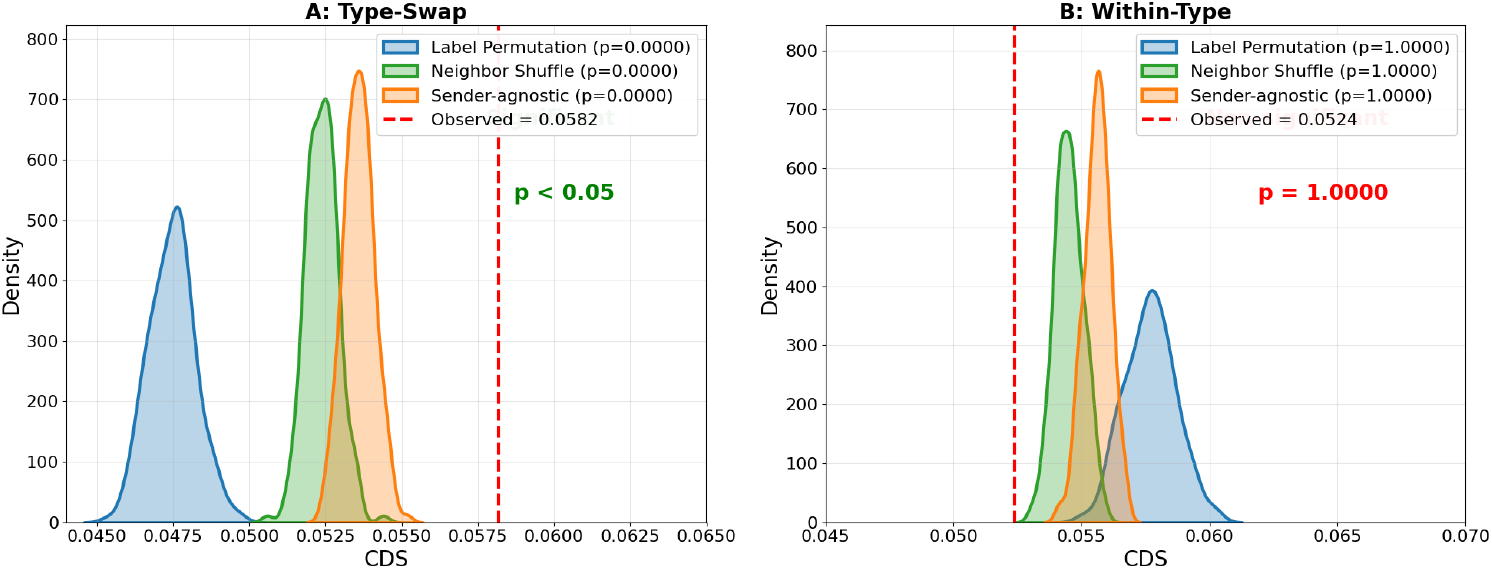
Null model comparison for Fibroblast→Macrophage. (A) Type-swap randomization. (B) Within-type constrained distribution.

Under the within-type shuffle operator (Fig. 5B), the observed CDS (0.0524) is within all corresponding null distributions (all *p* = 1.00). This indicates that the signal is driven primarily by Fibroblast type identity rather than intra-type state heterogeneity.

Bootstrap confidence intervals (reported above) further support stability across test cores. Per-core estimates are shown in Fig. 6.

**Fig. 6.**
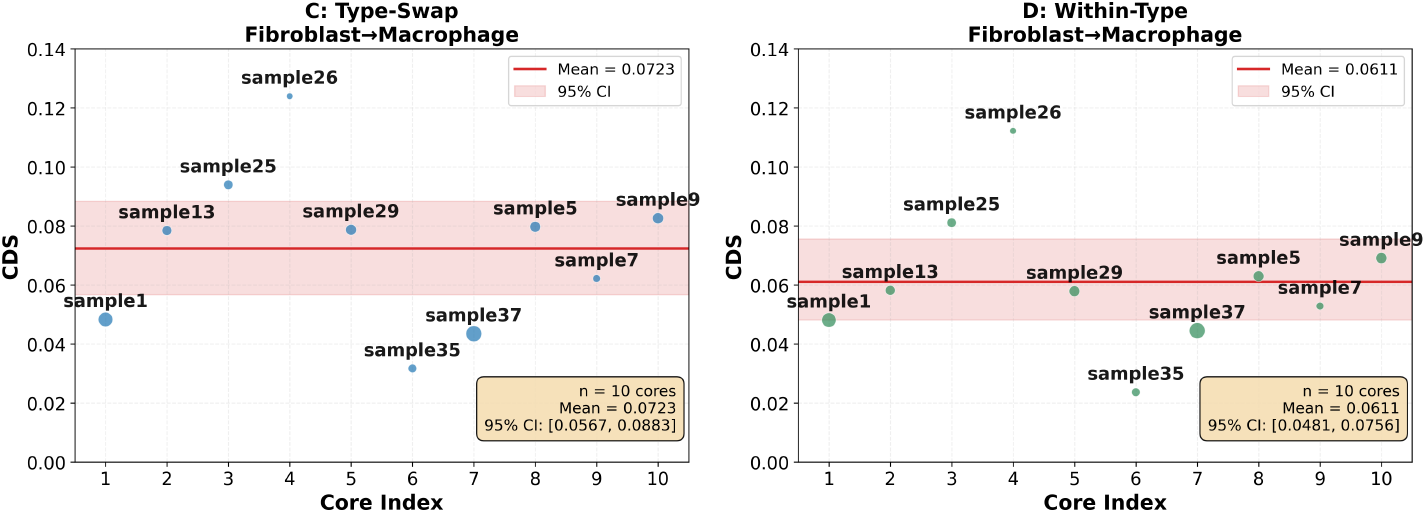
Cross-core CDS estimates for Fibroblast → Macrophage under type-swap (left) and within type (right) perturbations. Shaded region indicates bootstrap 95% CI.

### 3.6 Ligand–Receptor Concordance Validation

To evaluate biological concordance, we queried CellChatDB and computed ligand– receptor (LR) scores as the geometric mean of sender ligand and receiver receptor expression (mean expression *>* 0.05 in both cell types). Across 13 significant sender–receiver pairs, CDS values were strongly correlated with mean LR scores (*r* = 0.758, *p* = 0.0027), indicating alignment between counterfactually inferred directional influence and established signaling pathways. Fibroblast → Macrophage exhibited the highest LR score, consistent with its strong CDS.

### 3.7 Mechanistic Interpretability via Top Differentially Expressed Genes

To provide mechanistic detail for the observed directional influences, we identified, for each significant sender–receiver pair (FDR < 0.05), the three most up-regulated and three most down-regulated genes driving each of the top five impacted functional programs. The complete lists, including effect sizes (*Δ* values), are provided in Supplementary Tables S1a (up-regulated) and S1b (down-regulated).

For example, in the *Fibroblast* → *Macrophage* interaction (CDS = 0.0582, FDR < 0.05), the *Macrophage_Activation* program is driven by up-regulation of *CD68* (*Δ* = +0.011), *C1QC* (*Δ* = +0.008) and *FCGR3A* (*Δ* = +0.005), while the *EMT* program shows down-regulation of *POSTN* (*Δ* = − 0.043), *COL6A1* (*Δ* = − 0.024) and *COL6A2* (*Δ* = − 0.023). These gene-level insights directly support the biological interpretability of the counterfactual framework. The full lists of up-regulated and down-regulated genes per program are presented in Tables 1 and 2, respectively.

**Table 1.** Top up-regulated genes per functional program for significant sender–receiver pairs.

| Pair | Functional Program | Top Up-regulated Genes (delta) |
| --- | --- | --- |
| Fibroblast→Macrophage | Angiogenesis | — |
| Fibroblast→Macrophage | EMT | ITGB1 (+0.041) |
| Fibroblast→Macrophage | Macrophage_Activation | CD68 (+0.011), C1QC (+0.008), FCGR3A (+0.005) |
| Fibroblast→Macrophage | TGFb_Signaling | PMEPA1 (+0.028) |
| Fibroblast→Macrophage | Macrophage_TAM | LGALS3 (+0.012), MS4A7 (+0.005) |
| Neutrophil→Macrophage | Angiogenesis | — |
| Neutrophil→Macrophage | EMT | ITGB1 (+0.067), VIM (+0.014), POSTN (+0.011) |
| Neutrophil→Macrophage | Macrophage_Activation | C1QC (+0.003), CD68 (+0.003), FCGR3A (+0.002) |
| Neutrophil→Macrophage | TGFb_Signaling | PMEPA1 (+0.020), SPARC (+0.013), POSTN (+0.011) |
| Neutrophil→Macrophage | Macrophage_TAM | POSTN (+0.011), LGALS3 (+0.005), VCAN (+0.003) |
| Tumor_EMT→Macrophage | TGFb_Signaling | SPARC (+0.101), PMEPA1 (+0.042), POSTN (+0.040) |
| Tumor_EMT→Macrophage | Angiogenesis | — |
| Tumor_EMT→Macrophage | Macrophage_Activation | — |
| Tumor_EMT→Macrophage | EMT | POSTN (+0.040), COL6A3 (+0.024), COL6A2 (+0.020) |
| Tumor_EMT→Macrophage | Macrophage_TAM | POSTN (+0.040), VCAN (+0.033) |
| Fibroblast→CD8_T | Angiogenesis | — |
| Fibroblast→CD8_T | TGFb_Signaling | PMEPA1 (+0.019) |
| Fibroblast→CD8_T | EMT | — |
| Fibroblast→CD8_T | Macrophage_TAM | LGALS3 (+0.008), PLAC8 (+0.006) |
| Fibroblast→CD8_T | Macrophage_Activation | CD68 (+0.006), C1QC (+0.005), FCGR3A (+0.004) |
| Tumor_LD→Macrophage | Angiogenesis | CXCL12 (+0.016) |
| Tumor_LD→Macrophage | Macrophage_Activation | CD68 (+0.023), C1QC (+0.022), AIF1 (+0.015) |
| Tumor_LD→Macrophage | EMT | VIM (+0.037), POSTN (+0.036), COL6A1 (+0.025) |
| Tumor_LD→Macrophage | TGFb_Signaling | SPARC (+0.055), POSTN (+0.036), COL6A1 (+0.025) |
| Tumor_LD→Macrophage | Macrophage_TAM | POSTN (+0.036), MS4A7 (+0.024), VCAN (+0.018) |

**Table 2.**
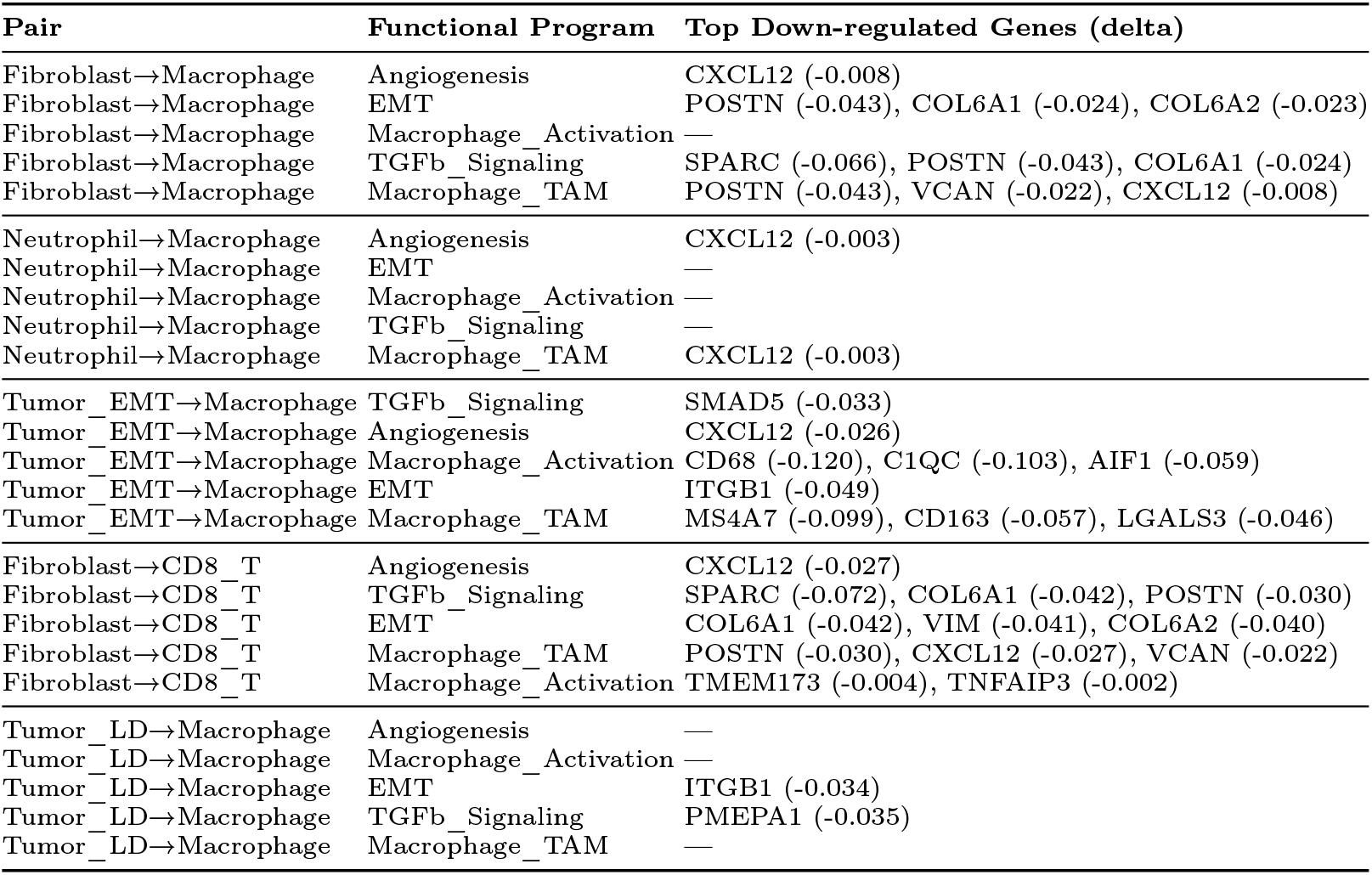
Top down-regulated genes per functional program for significant sender–receiver pairs.

### 3.8 Sensitivity Analysis

To assess robustness to neighborhood modeling choices, we performed a systematic sensitivity analysis for the *Fibroblast* → *Macrophage* interaction (the most stable pair with full core coverage). We varied (1) the number of neighbors *k* ∈ { 5, 10, 20, 30, 50 } and (2) the distance weighting scheme: exponential (default), uniform, inverse distance, and Gaussian (*σ* = 10 *µ*m). For each configuration, we recomputed the Counterfactual Directionality Score (CDS) with 500 bootstrap iterations across the 10 held-out test cores.

The results are summarized in Table 3. The weighting scheme coefficient of variation (CV) was 8.4%, indicating high robustness to the choice of distance weighting function. For *k*-sensitivity, we observed a CV of 17.7% with a clear monotonic decrease in CDS as *k* increased. This is an expected biological pattern: nearest neighbors exert the strongest influence on receiver cell state, while larger neighborhoods dilute this local signal. Importantly, across all configurations tested, CDS values remained significantly above null model expectations (*p <* 0.001), confirming that directional influence is detectable regardless of the specific neighborhood parameterization.

**Table 3.**
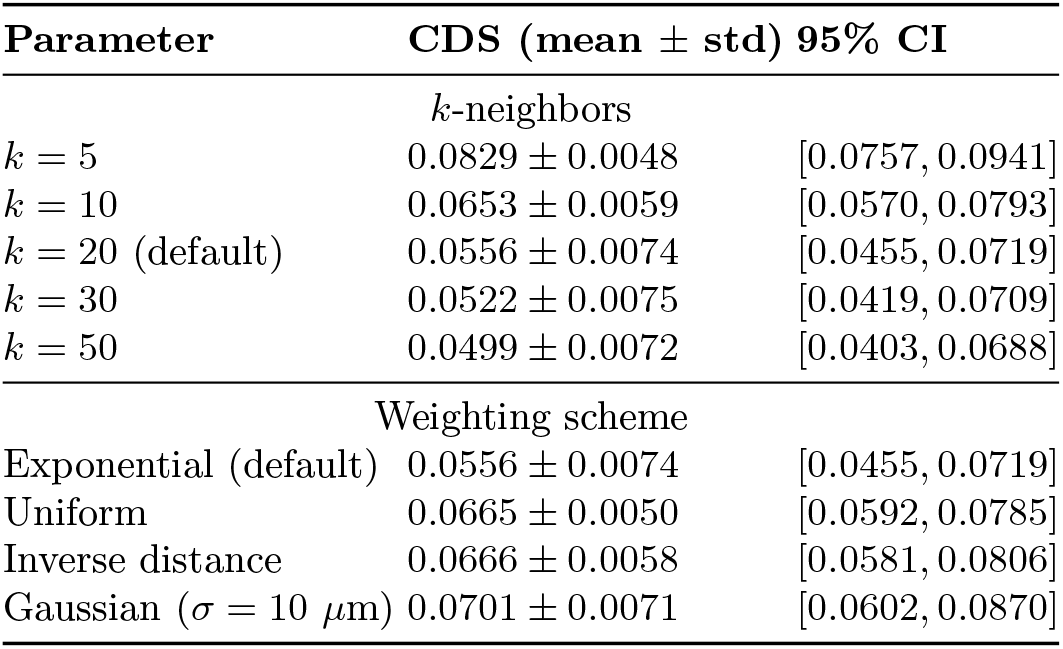
Sensitivity analysis for *Fibroblast*→*Macrophage* CDS.

Notably, the uniform and inverse distance schemes (which give higher weight to distant cells) produce slightly higher CDS values, while the exponential scheme (which prioritizes local neighbors) is more conservative. This further supports our choice of exponential weighting as a principled and conservative approach.

We selected *k* = 20 as the default because it balances sufficient spatial context to capture biologically relevant interactions while maintaining local specificity (CV stabilizes beyond *k* = 20), and it is consistent with standard practices in spatial transcriptomics neighborhood analysis.

## 4 Conclusion

We proposed a counterfactual framework for inferring directional cell–cell influence in spatial transcriptomics without predefined ligand–receptor priors. A Neighbor Influence Model predicts receiver cell state from spatial neighborhoods; controlled sender-type perturbations then define the CDS values. Applied to 10x Genomics Xenium based human cholangiocarcinoma TMAs, the framework revealed reproducible directional influences between tumor, immune, and stromal compartments, most prominently Tumor-EMT → Macrophage and Fibroblast → Macrophage. Three null models confirmed that observed effects are unlikely by chance, and ligand–receptor concordance analysis supported alignment with known biology (*r* = +0.758, *p* = 0.0027). These results demonstrate counterfactual testing as a statistically grounded alternative to correlation-based cell–cell communication analysis, applicable to large imaging transcriptomics datasets and clinical samples.

## Acknowledgment

This project was supported by the National Center for Advancing Translational Sciences (NCATS), National Institutes of Health, through Grant Award Number UM1TR004539. The content is solely the responsibility of the authors and does not necessarily represent the official views of the NIH.

## References

1. Browaeys, R., Saelens, W., Saeys, Y.: Nichenet: modeling intercellular communication by linking ligands to target genes. Nature Methods 17(2), 159–162 (2020). 10.1038/s41592-019-0667-5, https://doi.org/10.1038/s41592-019-0667-5

2. Cang, Z., Zhao, Y., Almet, A.A., Stabell, A., Ramos, R., Plikus, M.V., Atwood, S.X., Nie, Q.: Screening cell–cell communication in spatial transcriptomics via collective optimal transport. Nature Methods 20(2), 218–228 (2023). 10.1038/s41592-022-01728-4, https://doi.org/10.1038/s41592-022-01728-4

3. Chen, J.G., Chávez-Fuentes, J.C., O’Brien, M., Xu, J., Ruiz, E.C., Wang, W., Amin, I., Sheridan, J.P., Shin, S.C., Hasyagar, S.V., Sarfraz, I., Guckhool, P., Sistig, A., Jarzabek, V., Yuan, G.C., Dries, R.: Giotto suite: a multiscale and technology-agnostic spatial multiomics analysis ecosystem. Nature Methods 22(10), 2052–2064 (2025). 10.1038/s41592-025-02817-w, https://doi.org/10.1038/s41592-025-02817-w

4. Chen, T.Y., You, L., Hardillo, J.A.U., Chien, M.P.: Spatial transcriptomic technologies. Cells 12(16) (2023). 10.3390/cells12162042, https://www.mdpi.com/2073-4409/12/16/2042

5. Efremova, M., Vento-Tormo, M., Teichmann, S.A., Vento-Tormo, R.: Cell-phonedb: inferring cell–cell communication from combined expression of multi-subunit ligand–receptor complexes. Nature Protocols 15(4), 1484–1506 (2020). 10.1038/s41596-020-0292-x, https://doi.org/10.1038/s41596-020-0292-x

6. Jin, S., Plikus, M.V., Nie, Q.: Cellchat for systematic analysis of cell–cell communication from single-cell transcriptomics. Nature Protocols 20(1), 180–219 (2025). 10.1038/s41596-024-01045-4, https://doi.org/10.1038/s41596-024-01045-4

7. Liu, L., Chen, A., Li, Y., Mulder, J., Heyn, H., Xu, X.: Spatiotemporal omics for biology and medicine. Cell 187(17), 4488–4519 (2024). 10.1016/j.cell.2024.07.040, https://doi.org/10.1016/j.cell.2024.07.040

8. Liu, Z., Sun, D., Wang, C.: Evaluation of cell-cell interaction methods by integrating single-cell rna sequencing data with spatial information. Genome Biology 23(1), 218 (2022). 10.1186/s13059-022-02783-y, https://doi.org/10.1186/s13059-022-02783-y

9. Palla, G., Spitzer, H., Klein, M., Fischer, D., Schaar, A.C., Kuemmerle, L.B., Rybakov, S., Ibarra, I.L., Holmberg, O., Virshup, I., Lotfollahi, M., Richter, S., Theis, F.J.: Squidpy: a scalable framework for spatial omics analysis. Nature Methods 19(2), 171–178 (2022). 10.1038/s41592-021-01358-2, https://doi.org/10.1038/s41592-021-01358-2

10. Valihrach, L., Zucha, D., Abaffy, P., Kubista, M.: A practical guide to spatial transcriptomics. Molecular Aspects of Medicine 97, 101276 (2024). 10.1016/j.mam.2024.101276, https://www.sciencedirect.com/science/article/pii/S0098299724000359

11. Williams, C.G., Lee, H.J., Asatsuma, T., Vento-Tormo, R., Haque, A.: An introduction to spatial transcriptomics for biomedical research. Genome Medicine 14(1), 68 (2022). 10.1186/s13073-022-01075-1, https://doi.org/10.1186/s13073-022-01075-1

12. Xiao, X., Zhang, L., Zhao, H., Wang, Z.: Inferring spatial single-cell-level interactions through interpreting cell state and niche correlations learned by self-supervised graph transformer. Nature Machine Intelligence 8(1), 42–58 (2026). 10.1038/s42256-025-01161-0, https://doi.org/10.1038/s42256-025-01161-0

